# Ground tissue circuitry regulates organ complexity in cereal roots

**DOI:** 10.1101/2021.04.28.441892

**Authors:** Carlos Ortiz-Ramírez, Poliana Coqueiro Dias Araujo, Sanqiang Zhang, Edgar Demesa-Arevalo, Zhe Yan, Xiosa Xu, Ramin Rahni, Thomas R. Gingeras, David Jackson, Kimberly L. Gallagher, Kenneth D. Birnbaum

## Abstract

Most plant roots have multiple cortex layers that make up the bulk of the organ and play key roles in physiology, such as flood tolerance and symbiosis. However, little is known about the formation of cortical layers outside of the highly reduced anatomy of the model *Arabidopsis*. Here we use single-cell RNAseq to rapidly generate a cell resolution map of the maize root, revealing an alternative configuration of the tissue formative SHORT-ROOT (SHR) signaling pathway adjacent to an expanded cortex. We show that maize SHR protein is hypermobile, moving at least eight cell layers into the cortex. Higher-order SHR mutants in both maize and *Setaria* have reduced numbers of cortical layers, showing that the SHR pathway controls expansion of cortical tissue in grasses that sets up anatomical complexity and a host of key traits.

**One sentence summary:** Single-cell RNA-seq maps the maize root transcriptome uncovering a mechanism that regulates cortex layer number.

## Main text

Roots are radially symmetrical organs composed of three fundamental tissue types, the epidermis on the outside, the ground tissue at the middle, and a core of vascular elements plus pericycle that lie in a central cylinder known as the stele (*1*). The ground tissue is further divided into two different cell types, the endodermis and cortex, which are arranged as concentric layers around the stele. Variations in ground tissue patterning, particularly the number of cortex cell layers, are common across species and represent one of the defining features giving rise to interspecies root morphological diversity (*1*). This diversity allows plants to cope with biotic and abiotic stress and adapt to challenging environments. For example, roots with a multilayered cortex can develop mycorrhizal symbiotic associations, form specialized cortex-derived parenchyma for carbohydrate storage, and develop aerenchyma to avoid hypoxia in flood conditions (*2-4*). Therefore, an important ongoing question in root biology is how tissue patterning is adjusted to produce different root morphologies, and what are the underlying changes in the genetic networks controlling developmental differences in patterning among species.

A current limitation to answer this question is that our knowledge of radial patterning mechanisms in roots comes largely from the study of *Arabidopsis*, which possesses a simple root anatomy. In *Arabidopsis*, only two ground tissue layers develop in primary development, one endodermal and one cortical that originate from an asymmetrical cell division at or near the initials or stem cells (*5*). This division is controlled by the *SHORT-ROOT* (*SHR*)/*SCARECROW* (*SCR*) genetic pathway (*6, 7*). Mutants in either transcription factor develop a monolayered ground tissue. In addition, SHR mutants lack an endodermis, giving *SHR* a role in both tissue formation and cell identity (*8*).

Mechanistically, *SHR* functions as a mobile signal whose protein travels from the stele, where the gene is transcribed, into the surrounding endodermis, where it induces the expression of the downstream transcription factor *SCR* (*7*). The pathway then triggers the division that generates the cortex and endodermis layers (*8*). It has been suggested that additional movement of SHR from the stele further out into the cortex could cause extra cell divisions, giving rise to multiple cortex layers (*9*). However, the role of the pathway in a species with multiple cortex layers has not been functionally tested, and its involvement in cortical expansion has been unclear. For example, in *Cardamine hirsute*, a close relative of *A. thaliana*, the formation of multiple cortex layers appears to be independent of the SHR-SCR pathway, instead controlled by the activity of HD-ZIPIII proteins (*10*). Thus, the genes that control the elaboration of multiple cortex layers remain a central question. In addition, it is not known to what degree different layers of the morphologically similar cortex tissue are specialized in their function. One barrier to answering these questions is that high-resolution transcriptomics for comparative studies in species with multilayered cortex are lacking.

To address this gap, we first sought to produce a high-resolution spatial and temporal map of gene expression in a complex root that could provide clues to the genetic networks controlling morphological diversity in patterning. Therefore, we generated cell type-specific gene expression profiles using high-throughput single cell RNAseq (scRNAseq) to profile maize roots. Maize is a valuable model for comparative studies because its roots develop multilayered cortical tissues (8-9 cortex cell layers) within the root meristem, and it is amenable to protoplast generation, an essential step in plant scRNAseq (*11*). However, a major challenge of scRNAseq studies in species for which genomic resources are limited is the correct identification of cell types. Preliminary analysis showed that using homologs of *Arabidopsis* markers obtained by high-throughput cell sorting, did not provide a clear identification of morphologically homologous cell types in maize. This is likely because gene orthology is not always apparent and localization over such broad evolutionary distance is not well conserved. An alternative resource --an extensive set of tissue-specific markers for cell sorting, which was used to generate cell-identity marker lines in *Arabidopsis --* was not available in maize.

To overcome this problem, we first took advantage of the concentric arrangement of tissues in roots to develop a technique to fluorescently mark cell layers by dye penetrance (Dye Penetrance Labeling, DPL). In brief, a highly permeable dye (Syto 40 - blue) stains the entire root with low but detectable staining in stele (Fig. 1A), whereas a weakly permeable dye (Syto 81 - green) stains the outer tissue layers strongly, with a gradual drop in signal intensity towards the inner tissues (Fig.1A). This dual labeling was highly reproducible across roots and batches, allowing us to use the blue/green ratio gradient among the concentric layers of the root to separate inner from outer cell types using Fluorescence Activated Cell Sorting (FACS). We calibrated dye ratio to radial position by using DPL on a line expressing a fluorescent protein driven by the *SCR* promoter (*ZmSCR1*::RFP; Fig. 1B), which expresses in the endodermis, a middle layer of the root. RFP positive cells were used to calibrate a reference dye ratio for this middle layer, allowing demarcation of inner and outer tissues (Fig. S1A). We dissected root tips (5 mm from tip) and then rapidly digested their cell walls, sorting cells belonging to different tissue layers using their specific dye ratio. We also generated a set of whole meristem protoplasts vs intact root controls to filter out any effects of cell wall digestion. Importantly, digested and undigested controls clustered closely together and replicate samples yielded highly consistent profiles (Fig. S1B,C). To validate the DPL dataset, we compared our epidermal and stele layer sorts to a previous study that used mechanical separation of inner vs outer layers (*12*) and found an 80% agreement (Fig. 1D,E). In this manner, we developed a set of at least 170 markers for each radially arranged tissue (Fig. S1D). In addition, we obtained expression profiles of the root cap by dissection and quiescent center (QC), using FACS on a stable QC marker line, *ZmWOX5*::tagRFPt (Fig.S1E).

**Fig. 1.**
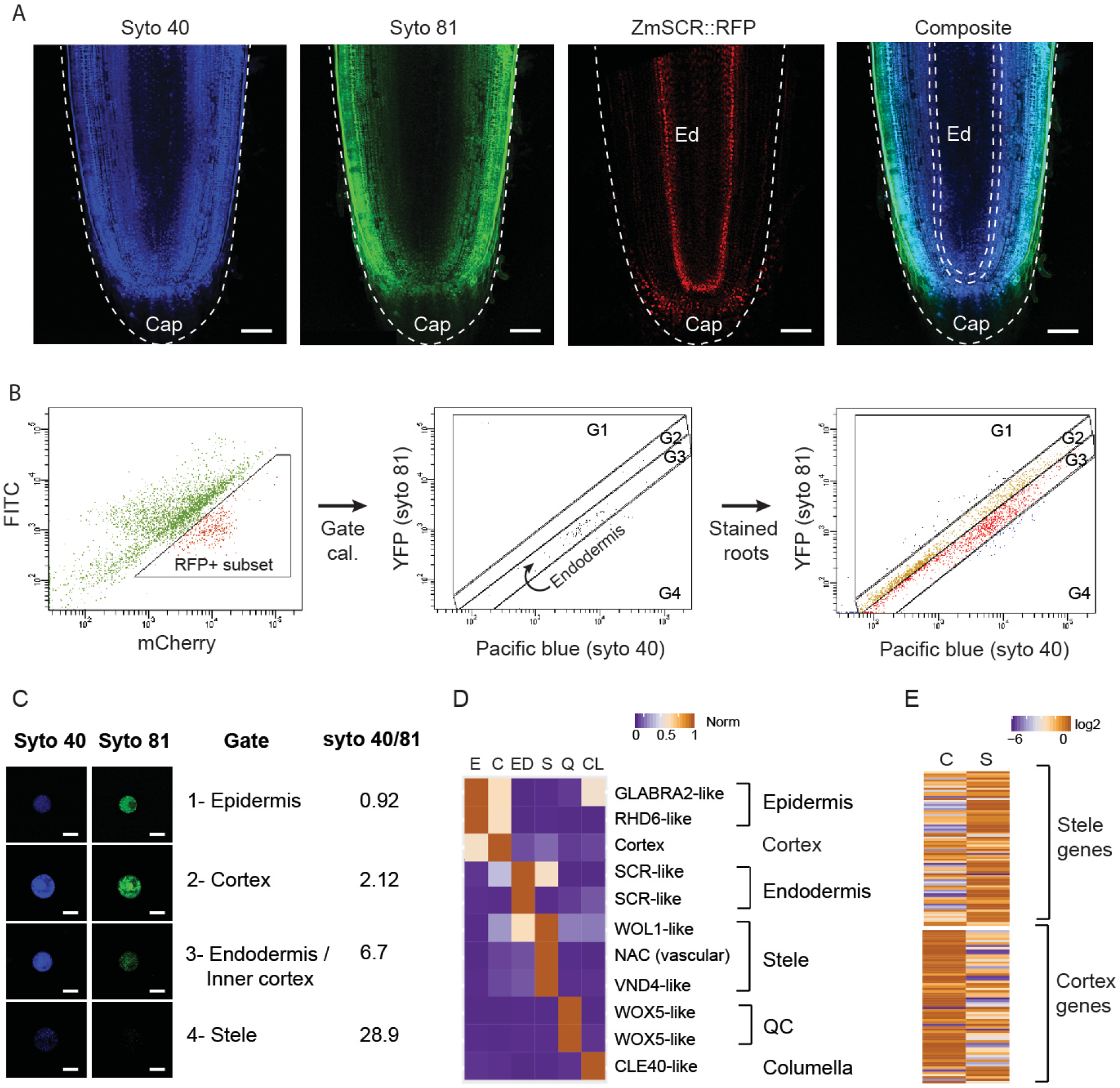
Dye Penetrance Labeling (DPL) and tissue transcriptome analysis in maize. (**A**) Representative images of a deeply penetrating dye (Syto 40), a superficially penetrating dye (Syto 81), the *ZmSCR*::tagRFPt marking endodermis, and a composite image of Syto 40 and Syto 81 staining, showing position of the endodermal (Ed) layers in dashed region. (**B**) Cell sorting gating strategy, showing the *ZmSCR*::tagRFPt population (left), backgated onto a YFP vs Pacific Blue plot with RFP positive (second from left), and (third from left) the gated boundaries for endodermal, outside of endodermis (G1,G2), and inside of endodermis (G4). (**C**) Validation of ratio metric cell sorting strategy by collecting sorted cells from gates and quantifying fluorescence from microscopy images. (**D**) Validation of sorted cell RNA-seq profiles via analysis of known markers. (**E**) Global validation comparing sorted cells vs. mechanically dissected stele and cortex tissues, with heat map showing expression in sorted cortex vs. stele gates, categorized by previously determined stele and cortex markers. Scale bars are 100 µm in (A) and 15 µm in (C).

To generate a single cell resolution map of the maize root meristem, we then dissected root tips from 7 day old wild-type B73 maize seedlings and enzymatically digested their cell walls, as above. We then used the cells to prepare single cell cDNA libraries using the 10x Genomics Chromium platform. A total of 4,324 high quality cells were sequenced in three different batches with a median of 15,254 UMIs/cell and 3,929 detected genes/cell. A total of 17 cell clusters were defined and visualized in two dimensions in Seurat using the uniform manifold approximation and projection (UMAP) method (*13*). To quantify cell identity and classify clusters using the DPL markers, we applied the Index of Cell Identity (ICI) algorithm (*14*), which generates a cell identity score based on the mean expression of a predefined marker gene set, in this case, from FACS isolated tissues (Fig. 2A & Fig. S2). Overall, the technique allowed us to identify, with high confidence, 16 of the 17 UMAP clusters, providing a detailed spatial map of transcripts in highly specific tissues of the maize root (Fig 2B).

**Figure 2.**
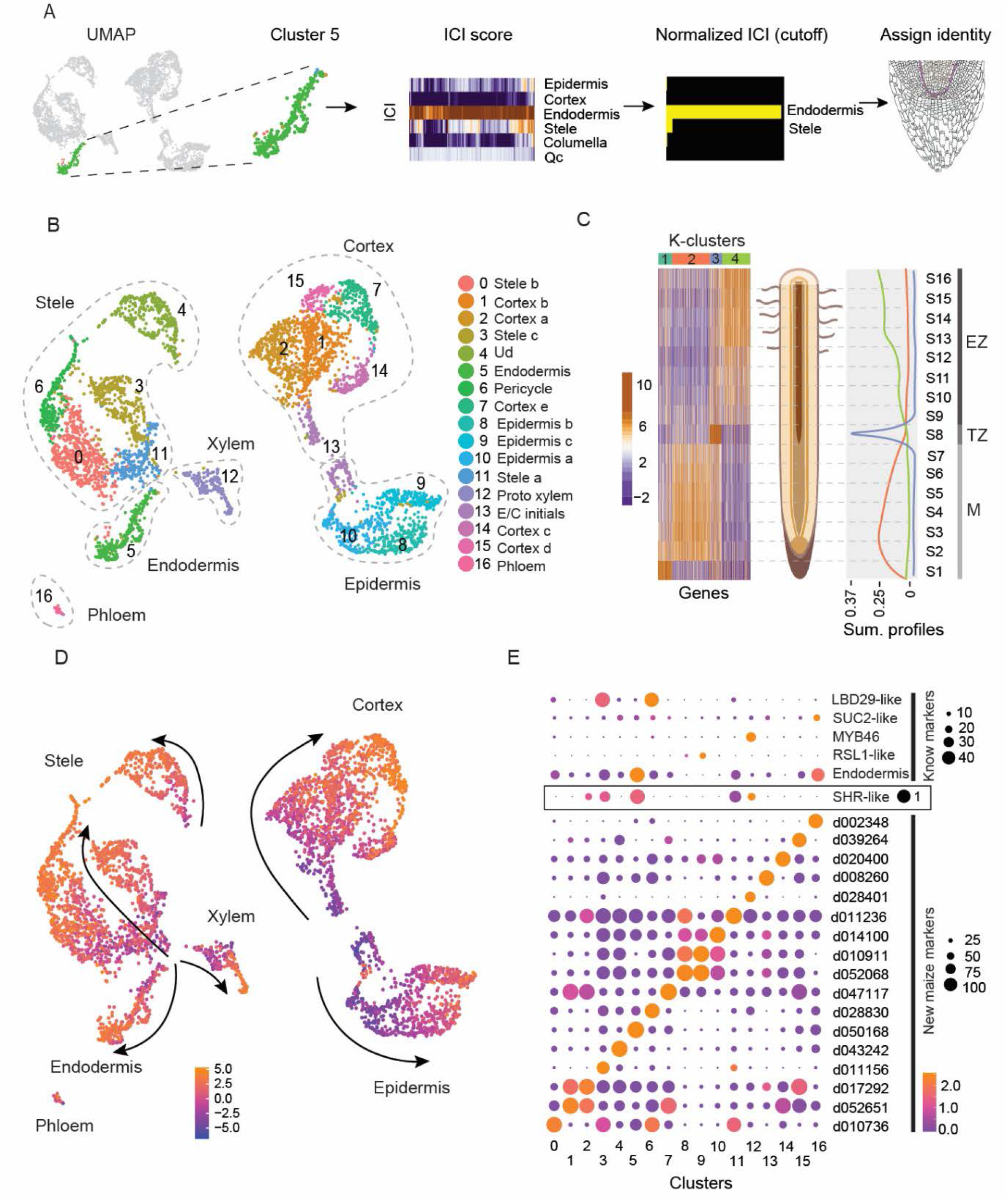
Single-cell RNA-seq spatial and temporal transcriptome maps of the maize meristem. (**A**) The ICI method of examining cells from a specific cluster in the UMAP analysis (e.g., Cluster 5), scoring the identity of each cell using tissue-specific markers obtained from DPL in Fig. 1, normalizing the scores by false discovery rate (see Methods). (**B**) Cluster identities determined by rough mapping (using ICI) and fine mapping using cell-specific markers. (**C**) Heat map of highly variant genes along a longitudinal axis of the root meristem, using thin sections of the root. Development patterns show transcripts/markers that peak in the early meristem (M), transition zone (TZ), and elongation zone (EZ). (**D**) Trajectories of developmental “pseudo-time” in each cell cluster mapped onto the same UMAP depicted in B, where a differentiation score is calculated as a log2 ratio of all EZ/M markers identified in C. (**E**) Validation of single-cell RNA-seq cluster calls with known markers (top). New markers for each cluster are shown at the bottom.

The high-resolution cellular map of the meristem showed multiple cell type subclusters within the stele and, interestingly, within the cortex as well, suggesting cellular specialization within that tissue’s multiple layers. However, because root cells differentiate as they transition away from the root tip, the possibility remained that some subclusters merely represented different maturation states of a single cell type. To distinguish groups formed by distinct identity rather than differentiation state, we further generated a set of cell maturation marker transcripts by dissecting 16 longitudinal root slices that together comprised the meristematic, transition, and elongation zones, and subjected the samples to RNAseq analysis (Fig. 2 C). Using K-means clustering, we identified three main expression programs: early differentiation transcripts (high expression on the meristem and gradual decrease towards the transition zone), transition zone transcripts (specific to the mid-maturation point), and late differentiation transcripts (low expression in the meristematic zone and gradual increase towards the elongation zone). We then generated a cellular differentiation score to label the differentiation status of each cell, resolving developmental trajectories of cells in our high-resolution map of cell identities (Fig. 2 D).

We found that, in a few cases, the state of cell maturation is indeed the main factor influencing grouping of cells into subclusters. Six clusters had the same identity as immediately adjacent clusters but representing a different state of maturation. However, the majority of subclusters were composed of cells with a wide range of differentiation states, showing that the grouping in most cases represented distinct cell identities. Importantly, while one cortex subcluster appeared to be a precursor state of mature cortex (cluster 1), our analysis confirmed the existence of at least four distinct cortex subtypes (Fig. 2B, clusters 2,1,14,15). Furthermore, using the ROC algorithm in Seurat, which identified 1,395 differentially expressed genes (DEGs) across all clusters, we found 446 transcripts that mark some subset of the four different cortical subtypes (Fig. 2E). Thus, we provide quantitative evidence for the sub specialization of cortex that underlies expansion of root complexity, providing strong evidence for cortical cell diversification.

A key question is what signaling mechanisms allow maize to form the extra layers that permit cortex sub specialization. We observed that a short list of functional markers with a role in patterning or cell identity in *Arabidopsis* had conserved localization in homologous tissues in maize (e.g., *SUC2* (phloem), *MYB46* (xylem), *RSL1* (epidermis), and *LBD29* (stele)). However, the localization of the core patterning gene *SHR* was surprising, as single cell data showed its expression was specific to the endodermis and not to the stele (Fig. 2E), where the *Arabidopsis* ortholog is expressed (*8*). All three maize paralogs of *SHR* (designated *ZmSHR1, ZmSHR2*, and *ZmSHR2-h*) showed the same endodermal enrichment in the profiles generated by single-cell and DPL/reporter analysis (Fig. 3A, B). We speculated that the expression of this mobile, division inducing transcription factor adjacent to the cortex could be related to a role in the expansion of that tissue.

**Figure 3.**
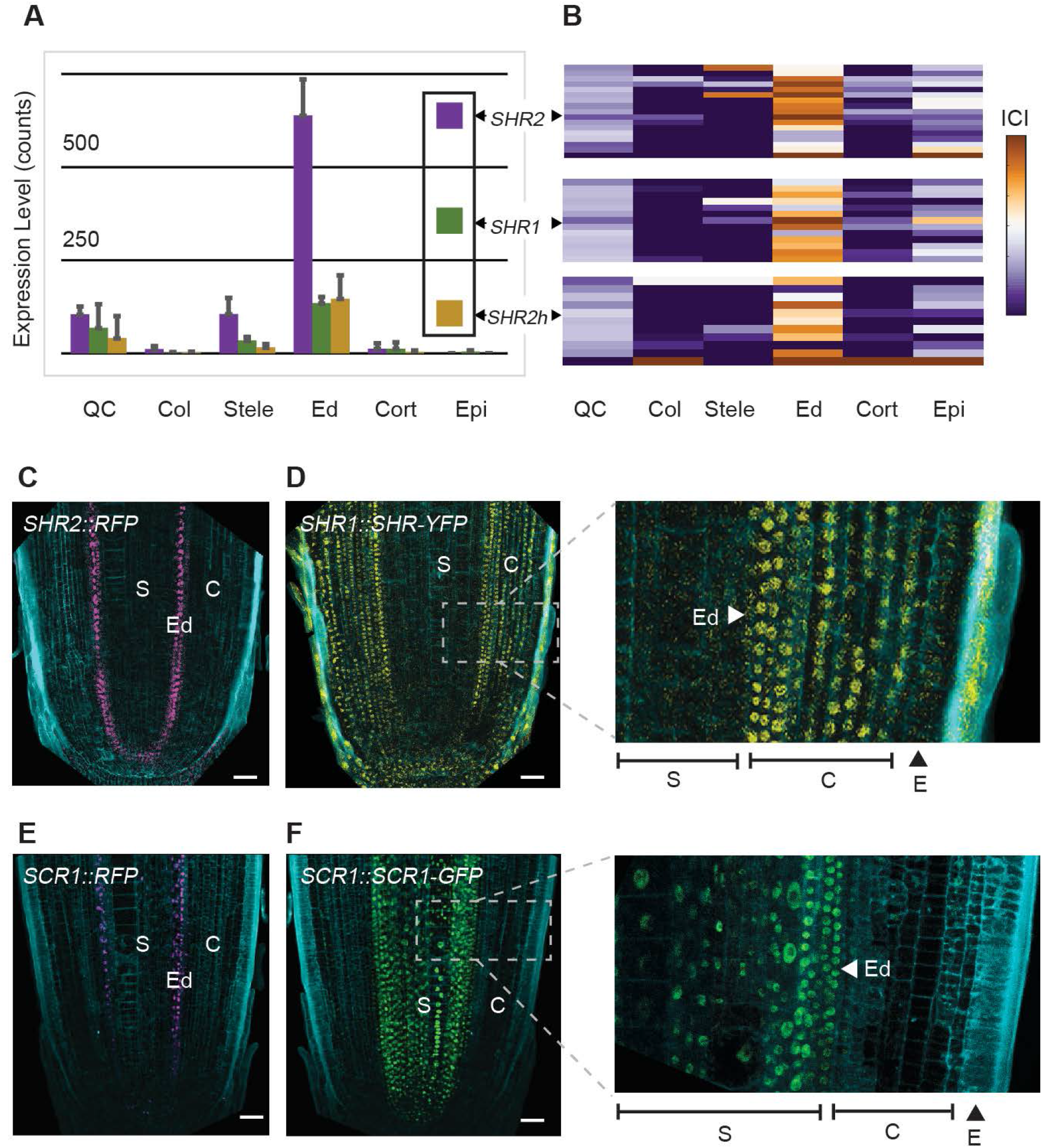
SHR and SCR expression in maize endodermis and differences between transcriptional and translational reporters. (**A**) Expression of the three *ZmSHR* paralogs (SHR1,2,2-h) as detected in sorted cells. Error bars are std based on three replicates. (**B**) ICI analysis of cells expressing either of the three *SHR* paralogs, identifying the SHR-expressing cells as having endodermal identity. (**C**) *ZmSHR2* transcriptional reporter. (**D**) *ZmSHR1* translational reporter, with inset showing enlarged view of boxed area and expression in all cell layers of cortex. (**E**) Transcriptional reporter for *ZmSCR1*. (**F**) Translational reporter for *Zm*SCR1 with inset showing enlarged view of boxed area and expression in many layers of the stele. Scale bars are 50 µm.

To confirm SHR transcript localization, we used 1.4 kb upstream and downstream regions of *ZmSHR2* to drive the expression of a nuclear-localized TagRFPt. Confocal images of root longitudinal sections from *ZmSHR1*::TagRFPt lines showed signal in the root endodermis in agreement with our dye sorted and single cell profiles (Fig. 3A,B,C). Notably, no signal was found in the stele, confirming that *SHR2* transcript localizes to different cell types in maize compared to *Arabidopsis*. Furthermore, given evidence that rice SHR proteins are hypermobile when expressed heterologously in *Arabidopsis* (8), we assessed whether maize SHR protein could move from the endodermis, where it is expressed, into the adjacent cortex. For this, we made a natively expressed protein reporter of ZmSHR1 fused to YFP (Zm*SHR1*::SHR1-YFP). Indeed, compared to endodermal localization of the *SHR1* found by dye sorting and single-cell RNA-seq, the maize SHR1 protein reporter was present in the cortex (Fig. 3D). Moreover, SHR1-YFP signal was not restricted to the immediately adjacent tissue layer as in *Arabidopsis* but was observed in all cortex layers, suggesting that the endogenous maize SHR1 protein moves through at least 8 cortex layers (Fig. 3D).

SHR’s role in promoting division in *Arabidopsis* works through direct interaction with SCR (*15*). Therefore, we also generated maize *SCR* reporter lines to determine colocalization with *SHR*. As noted in our DPL calibration, nuclear localized RFP expressed from the *ZmSCR* promoter revealed a clear signal in the root endodermis, showing *SCR* transcript localization is conserved in *Arabidopsis* and maize (Fig. 3E). However, natively expressed ZmSCR-GFP protein fusions showed a strong signal in the stele, suggesting that SCR protein in maize moves from cell-to-cell in the opposite direction to SHR (Fig. 3F). This shows that SHR and SCR colocalize only in the endodermis and not in the extra cortical layers, while SCR protein has an additional domain in the stele. A SCR translational reporter in a second monocot, *Setaria viridis* (Setaria), showed the same localization in the stele, further corroborating the divergent localization of SCR protein in monocots and suggesting SHR functions independently of SCR in the cortex (Fig. S3).

The model that implicates SHR in cortical expansion posits that increased outward movement of the protein could trigger periclinal cell divisions giving rise to extra ground tissue layers (*9*). To test the model, we targeted the three different maize SHR paralogs to generate loss-of-function mutants in maize using CRISPR-Cas9. We recovered mutants in two of the genes (*ZmSHR2* and *ZmSHR2-h*), including the most highly expressed paralog. Single mutants in *Zmshr2* and *Zmshr2-h* had no apparent difference in their root anatomy compared to wild type siblings. However, *Zmshr2/2-h* double mutants showed a significant reduction in the number of cortical layers: with most roots having 7 layers compared to 8 to 9 layers in wild type (Fig 4A,B,C,D). Mutants in the single *SHR* gene in *Arabidopsis* lack an endodermal layer. However, staining for suberin and morphological analysis showed that *Zmshr2/2-h* roots still developed an endodermis (Fig. S4). We posit that the remaining functional *ZmSHR1* gene in the *Zmshr2/2-h* background may still enable specification of endodermal identity. Alternatively, ZmSCR1 (and ZmSCR1-h) may be the primary factors in the specification of the maize endodermis (*16*). Overall, our results suggest that *SHR* function in maize is necessary for the full expansion of cortical identity.

**Fig. 4.**
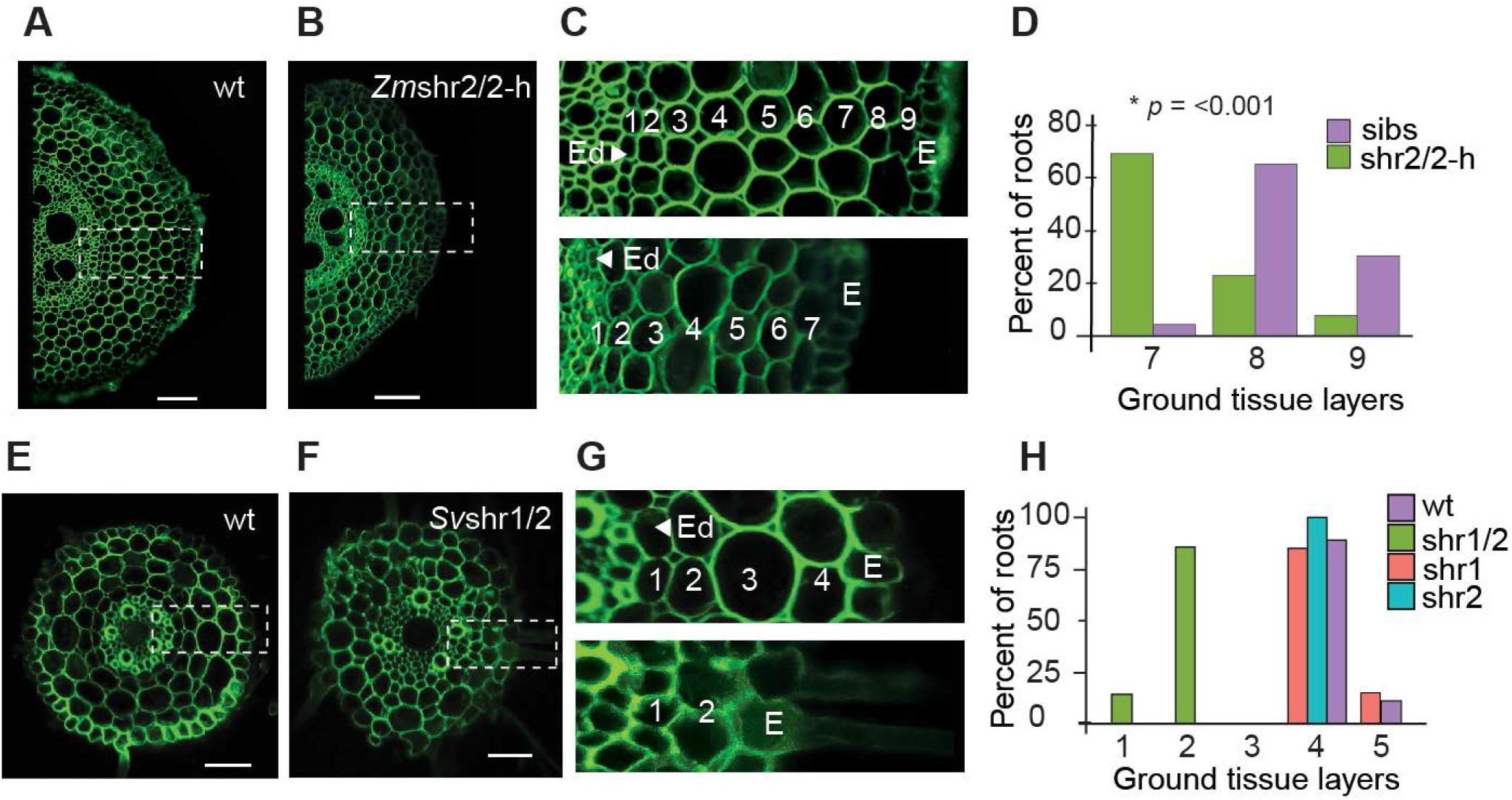
Cortical cell layer analysis in wild type and shr mutants in monocots. (**A**,**B**) Representative maize root cross-sections of wild type in A vs. *Zm*shr2/2-h double homozygous mutant in B. (**C**) Enlarged and labelled regions from dashed boxes in A and B showing stelar (S), endodermal (Ed), and cortical layers of wild type (top) and *Zm*shr2/2-h mutant (bottom). (**D**) Quantification of the cortical cell layers in wild type and het sibs (n=23) vs. *Zm*shr2/2-h mutant (n=13). *P* value was estimated with Mann-Whitney Rank Sum Test (**E**,**F**) Representative cross sections of *Setaria* wild type in E and *Sv*shr1/2 mutants in F. (**G**) Enlarged and labelled regions from dashed boxes in E and F showing stelar (S), endodermal (Ed), and cortical layers of wild type (top) and *Sv*shr1/2 mutant (bottom). (**H**) Quantification of cortical layers in *Setaria* wild type (n = 9), *Svshr1* single (n = 7), *Svshr2* single (n = 6) and double mutants (n =7). Scale bars are 100 µm in (A), (C), and 50 µm in (E), (F).

We sought to validate the monocot *SHR* phenotype with a more severe loss of function by testing its role in *Setaria viridis*, a close relative of maize. In Setaria, we were able to generate loss of function mutants in the two *SHR* orthologs using CRISPR-Cas9 (Fig. S5). One Setaria SHR mutant, Sv*shr2*, showed a slight reduction in cortical layers, while a single mutant in the second, Sv*shr1*, showed no phenotype. However, double mutants showed a dramatic reduction in the number of ground tissue layers: 1-2 layers compared to 4-5 layers in wild type siblings (Fig. 4E, F, G & H). These results corroborate the role of *SHR* in controlling the expansion of cortical layers in two monocots.

We note that the extra cortical divisions mediated by SHR could function through direct interaction with SCR by mediating successive divisions of the cortex-endodermal split near the stem cell niche, where the two proteins overlapped in maize. Alternatively, SHR hypermobility could lead to divisions directly in the cortical layers independently of direct interaction with SCR. At present, we cannot distinguish between these two models.

Notably, the results show that SHR has a role in monocots controlling the expansion of cortex, which sets up many key traits for environmental acclimation. This illustrates how a divergent role of a key patterning gene leads to a major difference in organ patterning. Furthermore, the results show that rapid transcriptome mapping using single cell dissection can provide insights into the mechanisms that mediate anatomical diversity. The use of dye labeling to generate a scaffold locational map together with scRNAseq now provides a detailed maize root tissue map that can be used as a reference map in monocots.

## Author contributions

C.O.R. performed all transcriptomic experiments, expression analysis, and mutant analysis. C.O.R. and K.D.B. designed the experiments and wrote the manuscript. C.O.R., K.D.B., D.J., T.R.G., and K.L.G conceived the project and guided the experiments. P.C.D.A. and S.Z. assisted in mutant analysis and marker analysis. E.D.A. and X.X. generated transcriptional reporters in maize. Z.Y. generated the translational reporters in maize and the Setaria mutants. R.R. designed and carried out microscopy protocols.

## Data materials and availability

All raw and processed/normalized expression data is available through the Gene Expression Omnibus under the SuperSeries accession GSE172302.

## Materials and Methods

### Accession numbers maize

SHR:

*Zm*00001d029607 (*Zm*SHR1)

*Zm*00001d021973 (*Zm*SHR2)

*Zm*00001d006721 (*Zm*SHR2-h)

SCR:

*Zm*00001d052380 (*Zm*SCR1)

*Zm*00001d005029 (*Zm*SCR2)

WOX5:

Zm00001d042821

(ZmWOX5B)

### Accession numbers Setaria

SHR

Sevir.9G361300 (SvSHR1)

Sevir2G383300 (SvSHR2)

SCR

Sevir8G008100 (SvSCR1)

### Plant growth Conditions

For protoplast generation and collection of root slices, maize seeds were incubated in 3% (v/v) sodium hypochlorite for 8 min. Seeds were then transferred to a sheet of wetted heavy-weight germination paper, rolled and covered with aluminum foil to prevent roots from exposure to direct light. Rolled paper was placed in a 2-gallon plastic container and kept inside a Percival high light chamber for 7 days with a dark-light cycle of 16 hr light at 27°C and 8hr dark at 24ºC and 50% humidity (*17*). For plant propgation and crosses, maize plants were grown inside a greenhouse with controlled light and temperature under a 16 hr light at 28°C and 8hr dark at 22°C for 3 months. *Setaria* seeds were sterilized as described before and germinated on square plates containing 0.5X Murashige and Skoog (MS) medium. Plates were kept in a Percival high light chamber for 14 days with a dark-light cycle of 16 hr light at 27°C and 8hr dark at 24°C and 50% humidity until collection of root tissue for microscopic analyses. Setaria plants used for seed bulking and genetic crosses were grown in a walk-in chamber exposed to the same light/dark cycle and temperature as described above.

### Generation of maize reporter lines

*Zm*SHR2, *Zm*SCR1 and *Zm*WOX5 transcriptional reporter lines were constructed using the Multisite GATEWAY (Invitrogen) recombination system using the pTF101 plasmid as a destination vector, all regulatory sequences were driving the expression of TagRFPt protein fused to four copies of the Nuclear Localization Signal (NLS) of VirD2 from *Agrobacterium tumefasciens* (*18*). For *Zm*SHR2, 1,405 bp of the upstream region and 1,445 bp of the downstream region were cloned into the reporter construct. For *Zm*SCR1, 1,704 bp of the upstream sequence and 2,054 bp of the downstream sequence were cloned into the construct. Finally, for *Zm*WOX5, 3,705 bp of upstream sequence and 2,826 bp of downstream sequence were cloned into the reporter. All expression plasmids were transformed in *Agrobacterium tumefasciens* strain EHA101 for maize transformation. At least two independent lines were used to verify the expression patterns.

For the ZmSHR translational fusion, the ZmSHR promoter (3,118 bp upstream from zmSHR), CDS and YFP were cloned and assembled into pTF101.1 destination vector via BamHI+NcoI, NcoI+SbfI and SbfI+AflII cloning site respectively. For the ZmSCR translational fusion, the ZmSCR promoter (3,261 bp upstream from zmSCR), CDS and GFP were cloned and assembled into the pTF101.1 destination vector via BamHI+BlpI, BlpI+SbfI and SbfI+EcoRI cloning site respectively. All expression plasmids were transformed into *Agrobacterium tumefasciens* strain EHA101 and maize transformation was performed at the Plant Transformation Facility of Iowa State University.

### CRISPR-Cas9 mediated gene editing of maize SHR genes

Null alleles for *Zm*SHR2 and *Zm*SHR2-h were generated using the CRISPR-Cas9 system. Five sgRNAs were designed to simultaneously target both SHR genes using CRISPR-P 2.0 web-tool(*19*) based on B73 reference genome (GCAACACGTTCTGCACGCAG, GTCGAAGGACGTGTTCCGGT, GCATTTCCACTCGCACGGCG, GCACTCCAGCAGCAGCTGCG). The array of sgRNAs under the ZmU6 and OzU6 promoters were synthesized and cloned into a pGW-Cas9 construct and transformed into the maize Hi-II accession using *Agrobacterium tumefasciens* (*20*). Transformed plants were screened by PCR amplification of the genomic region containing the targeted site. Transformed plants were backcrossed once to B73 and then plants with edited alleles were crossed reciprocally to segregate both *Zm*SHR2 and *Zm*SHR2-h edited alleles under the same genetic background. All phenotypic assays were done between plants harboring null alleles and their segregating siblings.

### SCR translational fusion in Setaria

For SvSCR translational fusion, SvSCR promoter (2 kb upstream)+5UTR, and SviSCR CDS were assembled into modified PENTR 3C-YFP vector via SbfI+BamHI and BamHI+EcoRI cloning site, respectively. pSvSCR-SvSCR-YFP was cloned into modified PANIC 10A-Gateway destination vector using GATEWAY (Invitrogen) recombination system.

### CRISPR-Cas9 mediated editing of Setaria SHR genes

sgRNA were designed using CRISPR-P (svSHR1: CGTGGCCGAACGACGCCCAC; svSHR2: TCTGCTAGAGTGCGCTCGGG). The sgRNA of svSHR1 was assembled into pJG338 and the sgRNA of svSHR2 was assembled into pJG340 via the BaeI cloning site. The vectors pJG471 containing Ubi1::TaCas9 and pJG338 or pJG340 were assembled into the pRLG103 destination vector via the AarI cloning site. Stable *Setaria viridis* transformation with CRISPR-Cas9 vectors was performed as described previously (*21*). T0 and T1 plants were screened by PCR targeting CRISPR editing site of svSHR1 or svSHR2.

### Dye Penetrance Labeling (DPL)

Roots tips (5mm) from 7-day-old plantlets were placed on a tissue culture dish containing a solution of Syto 40 (1µM) and Syto 80 (0.6µM) in 3ml of diH_2_O containing L-cysteine (2.5 mM) for 40 min, shaking continuously at 60 rpm. Stained roots were washed at least 5 times in diH_2_O and transferred to protoplast generating solution (see below). In preliminary work, variation in staining was minimal from root to root, and modifying incubation time only changed fluorescent intensity but did not modified penetrance of either dye. Using the ratio of Syto40 to Syto 80 rather than absolute signal of either normalized for root-to-root variation.

### Protoplast Generation

Protoplasts were generated from primary and seminal roots as described previously (*17*). In brief, roots were cut above the meristem at approximately 4 to 5 mm from the root tip and placed in pretreatment solution containing L-cysteine for 40 min (0.75 g sorbitol, 62 µl of 1 M L-cysteine in 25 ml) to improve enzyme efficiency and cell wall digestion. When root tips were used for DPL, the pretreatment solution also contained Syto 40 and Syto 81 dyes and without sorbitol, as it decreases dye penetrance efficiency. Cell walls were digested for 90 min in an enzyme solution optimized for maize roots [1.2% cellulase “Onozuka” RS, 1.2% cellulase “Onozuka” R10, 0.4% macerozyme R-10 (all three Yakult Pharmaceutical Industry CO.), 0.36% pectolyase Y-23 (MP Biomedicals)]. Protoplast were then filtered through a 40 µm cell strainer and transferred to microcentrifuge tubes for centrifugation (3 min at 500 x g). Protoplast pellets were washed and resuspended in washing solution (0.4 M mannitol, 0.02 M MES, 0.02 M KCl, 0.01 M CaCl_2,_ 0.015 mM BSA) and used immediately for fluorescence-activated cell sorting or single-cell RNAseq.

### Cell Sorting and Tissue Profiles

A gating strategy was set up to delineate four gates, representing epidermis, cortex, endodermis and stele. First, we stained *Zm*SCR::RFP line using the DPL technique outlined above with Syto 81 and Syto 40. Protoplasts generated after staining were used to first identify RFP positive cells in a FITC vs. mCherry set of panels on a FACSAria II (Becton Dickinson). Those same cells were also visualized (backgated) in a YFP (for Syto 81) vs. Pacific Blue (for Syto 40) plot, allowing us to determine a gate that represented the narrow range of ratios that represented endodermal cells in YFP vs. Pacific Blue plot (G3). Using DPL and short digestion protocols that largely released epidermal protoplasts, we similarly determined the ratio range for epidermal cells (G1). The ratio range (Syto 80/Syto 40) between epidermal and endodermal cells was then labelled as the cortical layers (G2). The ratio range higher than the endodermal gate was labeled as stele (G4). A fifth sample representing the quiescent center (QC, with some xylem expression) was collected separately using a pWOX5::tagRFPt line and a similar gating strategy used before to detect the SCR::RFP line in the FITC vs. mCherry plot. For root cap isolation, we took advantage of the intrinsic properties of the cap, which is not amenable to enzymatic digestion. Root tips were thus digested with enzymatic solution for 90 min effectively removing cells of the root proper, and the remaining caps were inspected under a stereomicroscope and collected (30 caps for each replicate). All samples were subjected to the same RNA extraction and library preparation detailed below.

### Protoplast and Dye Staining Controls

Controls were generated to assess the effects of protoplast generation: For protoplast generation, approximately 30 maize plants were germinated in germination paper and grown for 7 days as described above. Primary and seminal roots tips (5mm) were harvested and split in half arbitrarily. Half of the roots were subjected to cell wall digestion as above, while the second half was processed for RNA extraction after manual dissection of the cap. Caps were removed to make the cell composition in each sample the same, since digested root preparations lack protoplasts from the cap.

### Root Slices

Roots were grown on germination paper as described above and cross sections of primary roots about 80-100 µm thick were harvested at 7 days after germination in the following manner: three roots tips (10 mm long) were cut and placed on top of a microscope slide covered by a rectangular piece of germination paper. Root tips were aligned parallel to each other under a stereoscope, taking the tip of the roots as a refence. Once aligned, the germination paper was bent around and placed on top of the root tips covering the last 5 mm (proximal part of the root). A small balance weight (300g) was placed on top the germination paper to hold the roots in position while cutting the cross sections. Cross sections were cut using a feather microscalpel (Electron microscopy sciences, cat. 72045-30) starting from the tip of the root to the proximal region. 16 equivalent slices were obtained from two roots (replicates) and collected in individual tubes containing RTL buffer (Qiagen MicroRNAeasy Kit). Samples were flash frozen by placing liquid nitrogen in the tubes. RNA extraction and library preparation were done as described below.

### Small Sample RNA-seq

RNA was extracted using the Qiagen MicroRNAeasy Kit. For dissected tissue samples (slices, protoplasting controls, and root cap), tissue samples were macerated in a 1.5 ml Eppendorf filled with RLT buffer and ground by hand with a plastic pestle after flash freezing with liquid nitrogen. For sorted cells, the positive flow stream was sorted directly into RLT buffer, after which point the Qiagen MicroRNAeasy extraction protocol was performed, excluding tissue homogenization. We performed cDNA construction and library amplification with Nugen OvationRNA Amplification System V2 (Tecan). Libraries were made using the Nugen Ovation Ultralow DR Multiplex System (Tecan). DNA fragmentation was performed using the Covaris S220. Short read sequences were generated on an Illumina NextSeq 500 and were aligned using HISAT2 to the maize B73 v4 reference genome. Counts were normalized between samples using DESeq2’s median of ratios method before analyzing for differentially expressed genes.

### Single cell RNA-seq

Protoplasts were prepared as described above. An aliquot was stained with flourescein diacetate (2µg/ml) for 3 min to determine viability before loading. Approximately 10,000 cells were loaded in a Single Cell A Chip (10x Genomics) per replicate. Three independent replicates were performed. Chips were loaded on a Chromium Controller (10x Genomics) to generate single-cell GEMs. Single-cell libraries were then prepared using the Chromium Single Cell 3’
s library kit, following manufacturer instructions. The quality of resulting DNA libraries was assessed with an Agilent TapeStation system. Library concentration was determined by quantitative PCR (qPCR) and sequenced with an Illumina NextSeq 550 platform using a 1×150 high-output configuration. Raw scRNAseq data was analyzed by Cell Ranger 2.2.0 (10x Genomics) to generate gene-cell matrices. Gene reads were aligned to maize B73 v4 reference genome.

### UMAP and ICI analysis

Replicates (3 independently generated single-cell libraries) were integrated and cells mapped using the Seurat package v3.0 (*13*) as follows: first, genes with counts in fewer than three cells were excluded from the analysis and their counts were removed. Second, any cell with fewer than 200 total UMI were also removed. Next, counts were log-normalized and the 2000 most variable genes were identified for each replicate using the “vst” method in Seurat. Next, we used the *FindIntegrationAnchors* function to identify anchors between the three datasets, using 20 dimensions. A new profile with an integrated expression matrix containing cells from all replicates was produced with the *IntegrateData* function. For dimensionality reduction, the integrated expression matrix was scaled (linear transformed) using the *ScaleData* function, and Principal Component analysis (PCA) performed. The top 20 principal components were selected. Cells were clustered using a K-nearest neighbor (KNN) graph, which is based on the Euclidean distance in PCA space. The *FindNeighbors* and *FindClusters* function with a resolution of 0.6. was applied. Next, non-linear dimensional reduction was performed using the UMAP algorithm with the top 20 PCs.

For single cell ICI analysis, high quality cells were selected according to default cutoffs in CellRanger (10x Genomics). Each cluster was then analyzed using the Index of Cell Identity approach (*14*) using 50 markers for each of the six tissues isolated using the dye penetrance technique (4 tissues) and the two markers used to sort cells, *Zm*SCR::RFP and *Zm*WOX5::RFP. Markers were selected by correlation to tissue specific pattern including replicates (e.g., 1 1 1 0 0 0 0 0 0 0 0 0 0 0 0), with a cutoff of r=0.85 using 50 highest correlated genes as markers for each tissue. The False Discovery Rate was calculated by randomizing the marker by tissue type matrix 1000 times and using the 95^th^ percentile highest score for a given tissue as a cutoff for a 5% FDR. The heatmap represents the log2 values of the observed ICI-95^th^ percentile cutoff, which we termed the normalized ICI in Fig. 1. The ICI score for cells expressing SHR1, 2, or 2-h in Fig. 3 was calculated similarly.

### Microscopy for marker and mutant Analysis

Quantification of ground tissue layers from wild type and mutant roots in maize and *Setaria* were performed by hand sectioning cross section “slices” of approximately 100 µm acquired at approximately 10 mm from the tip. Cross sections were flipped onto the flat surface of a slide, mounted in water, and imaged using a Leica SPE confocal microscope. Samples were excited at 405 nm generating an autofluorescent profile that outlined cell walls collected at 505-555 nm. Maize and *Setaria* fluorescent reporter lines were imaged by performing longitudinal cuts of primary and seminal roots as close as possible to median. As above, root sections were mounted in water. The RFP markers were imaged with a 561 nm laser using an mCherry filter setting, while GFP and YFP markers were imaged with a 488 laser using FITC and YFP filter settings on the Leica SPE.

### Suberin Staining

Anatomical characterization of wild type and mutant plants in maize was performed by first fixing roots in 50% ethanol. Hand sectioned “slices” were sampled in the late differentiation zone just below the epicotyl junction. Root slices were stained with fluorol yellow 088 stain (0.01% w/v, Sigma) in lactic acid (Sigma) in 70°C water bathing for 1 hour. The sections were then washed by deionized water and post-stained with 0.1% (w/v) toluidine blue O (Fisher scientific) for 2 min.

## Supplemental Figures

**Fig. S1.**
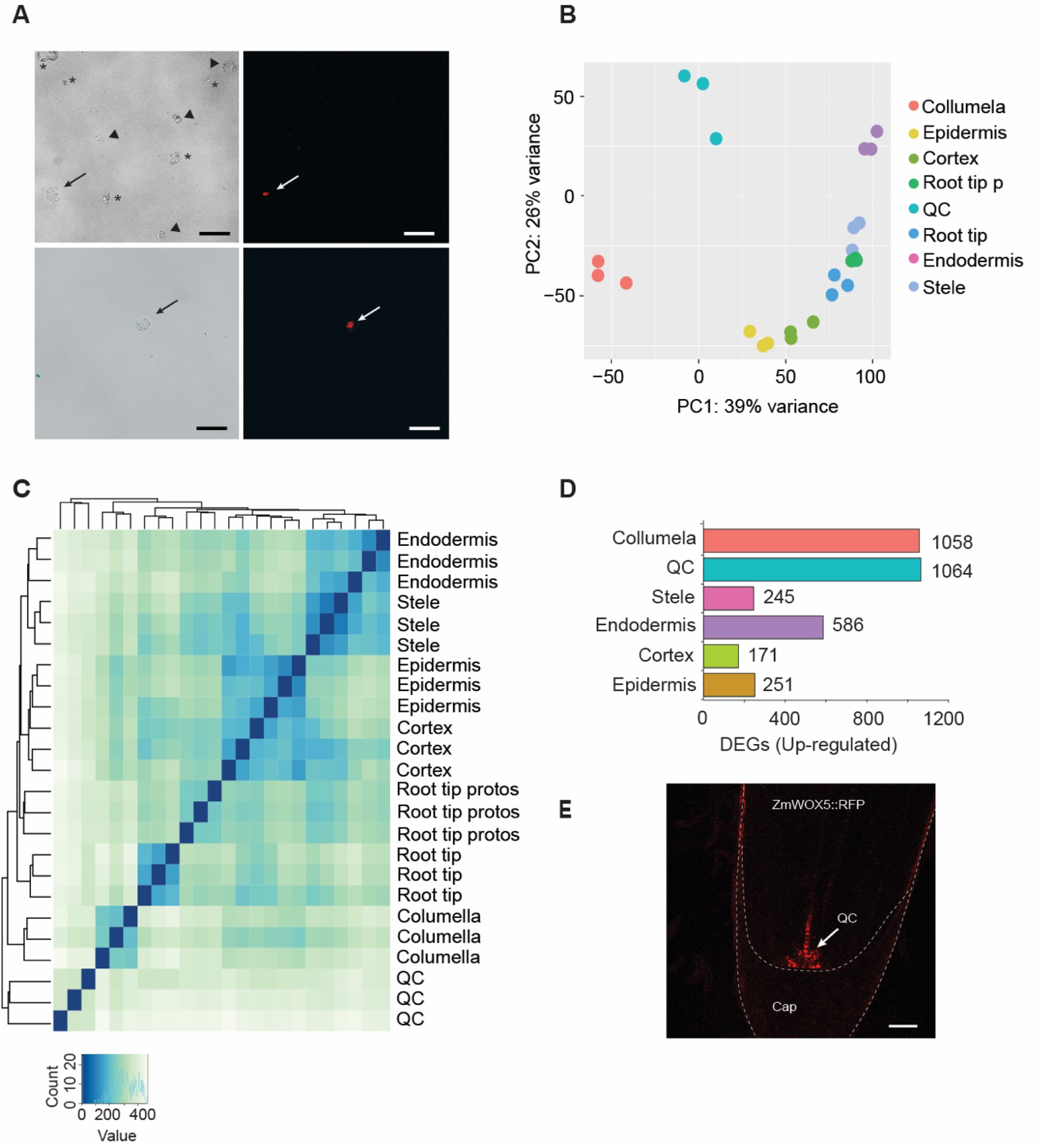
FACS sorting of RPF positive and dye-stained protoplasts and clustering analyses of tissue transcriptomes. **(A)** Bright field microscopy showing protoplasts (triangles) and debris (asterisks) from *ZmSCR*::tagRFP1 digested roots before (top left) and after (bottom left) FACS sorting. Fluorescence confocal microscopy confirmed that protoplasts expressing RFP were present (arrow, top right) and isolated (arrow, bottom right). **(B)** Principal Component Analysis (PCA) of tissue-specific gene expression patterns and whole roots vs. all root protoplast controls. **(C)** Hierarchical clustering dendrogram of tissues replicates showing sample similarity. **(D)** Number of upregulated differentially expressed genes. Root tip represents the undigested root meristem and root tip p/protos represents the same region of the root as a collection of protoplasts. Scale bars are 50 µm in A and 100 µm in E.

**Fig. S2.**
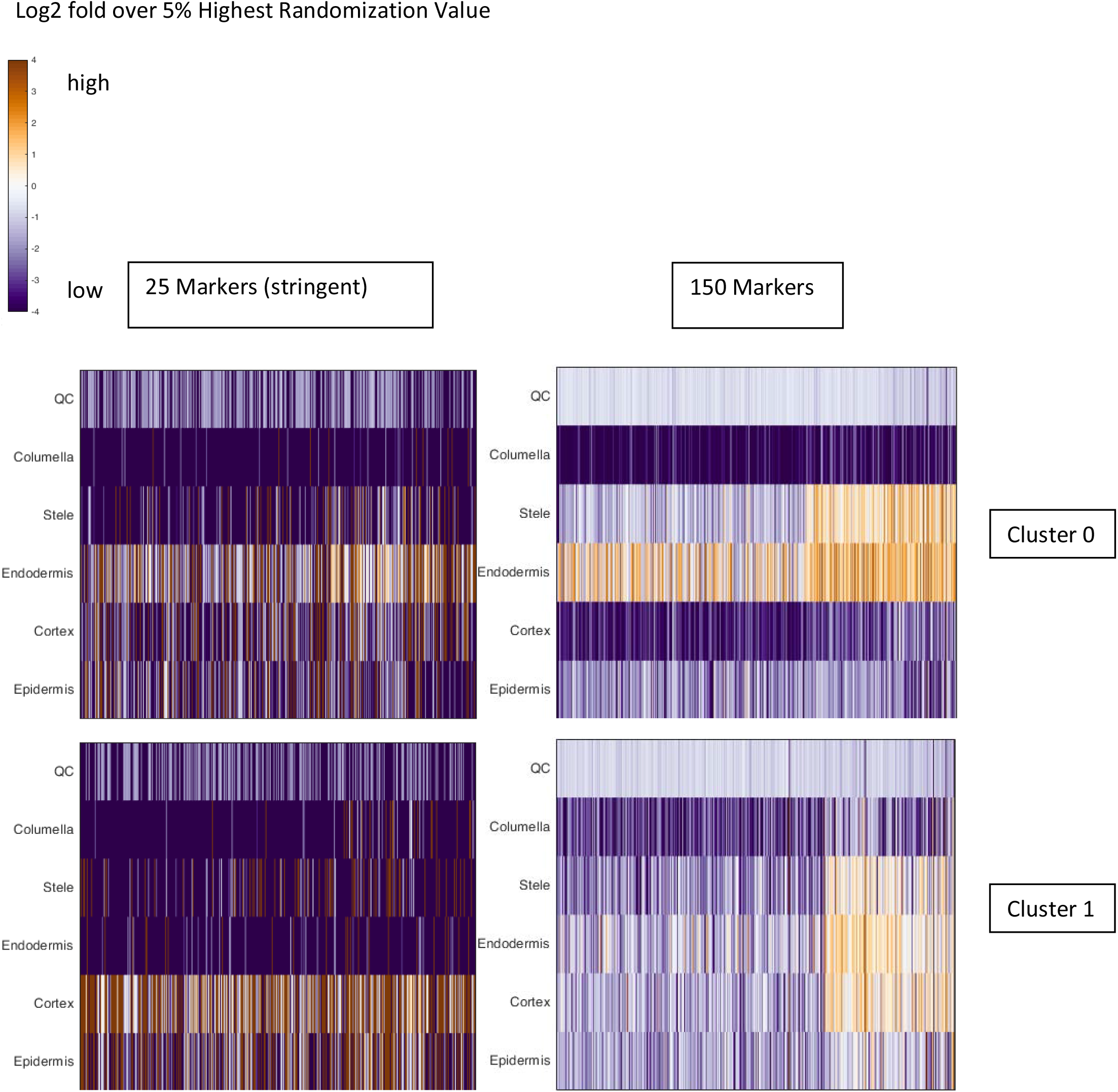
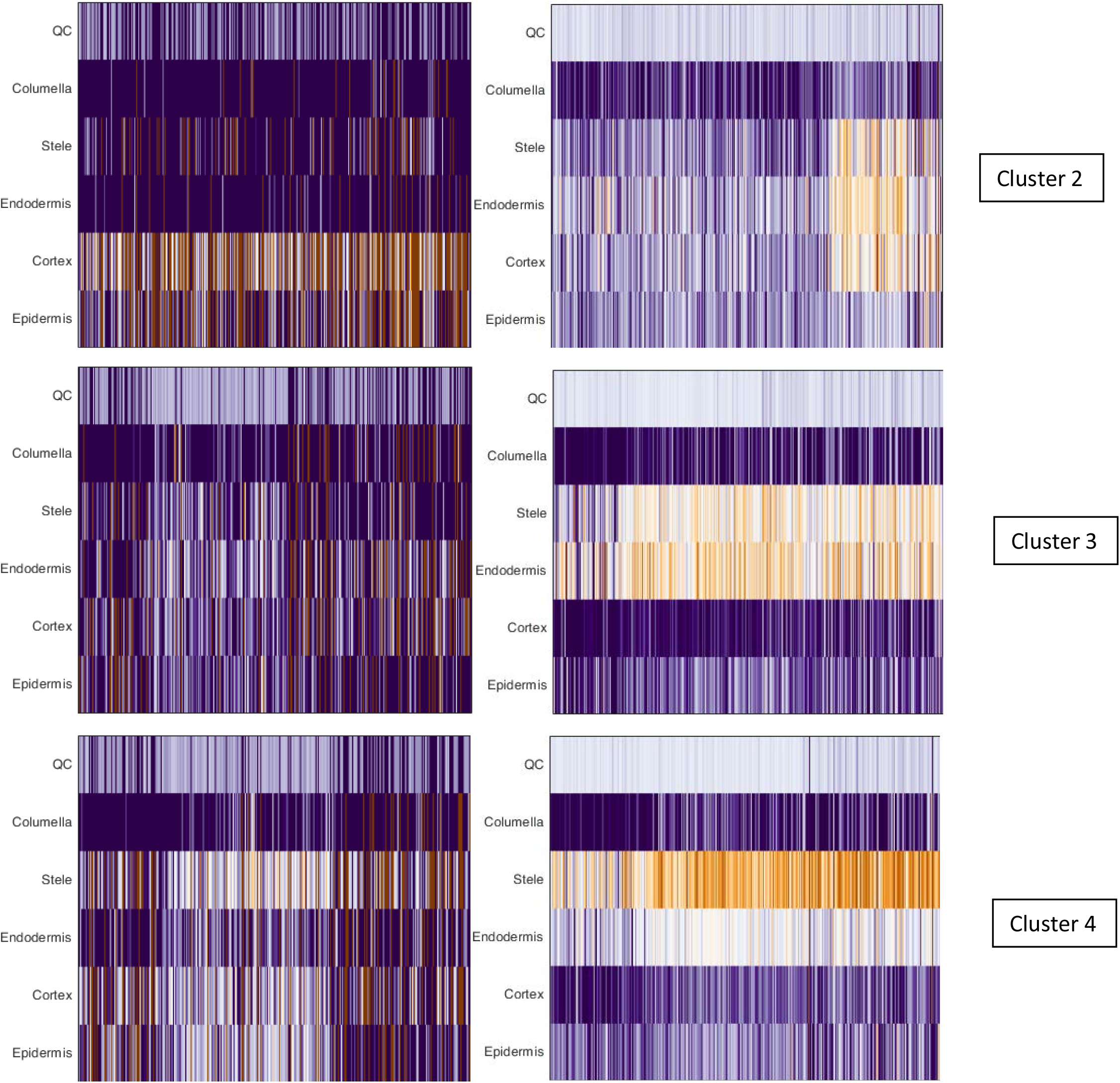
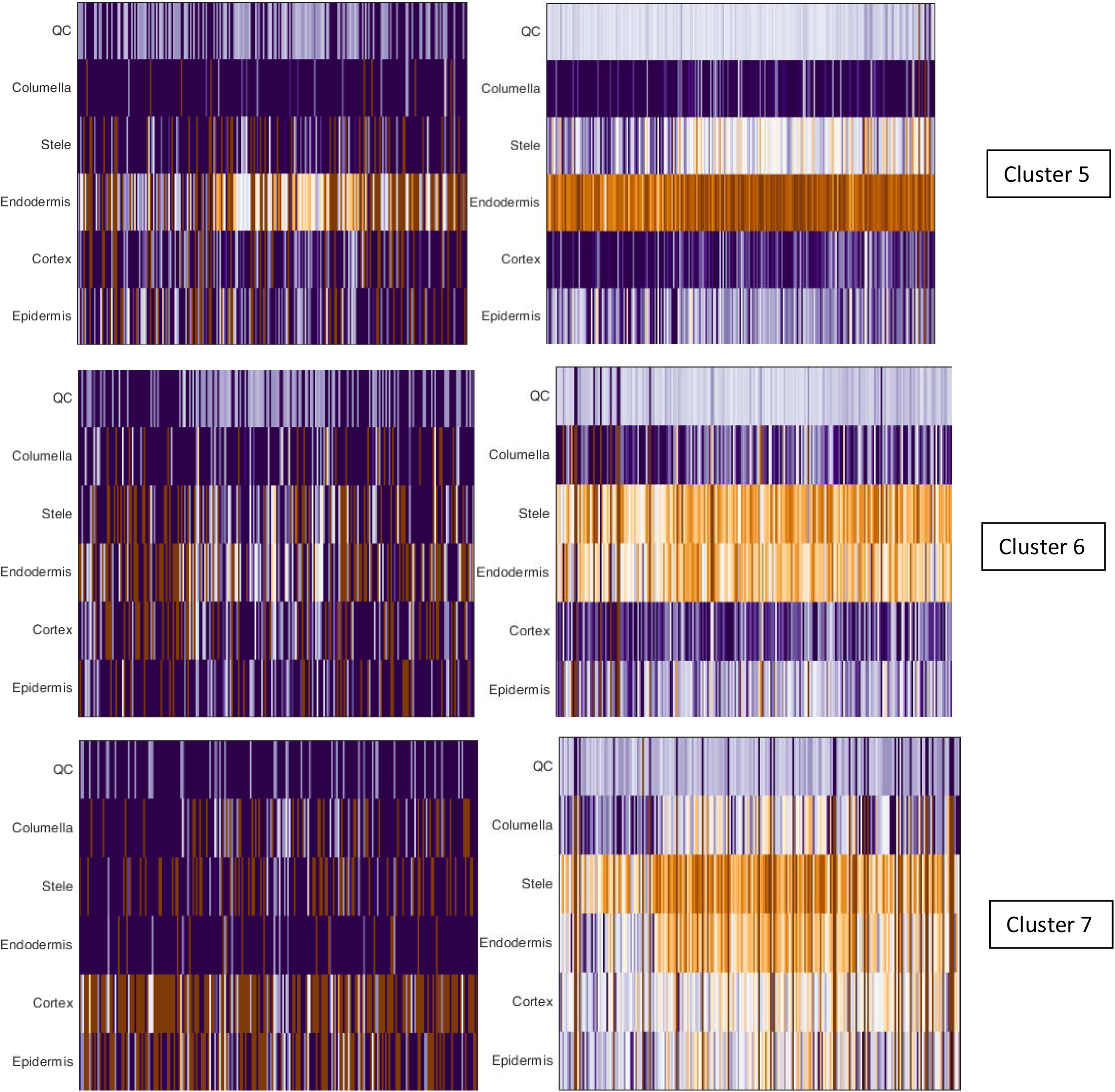
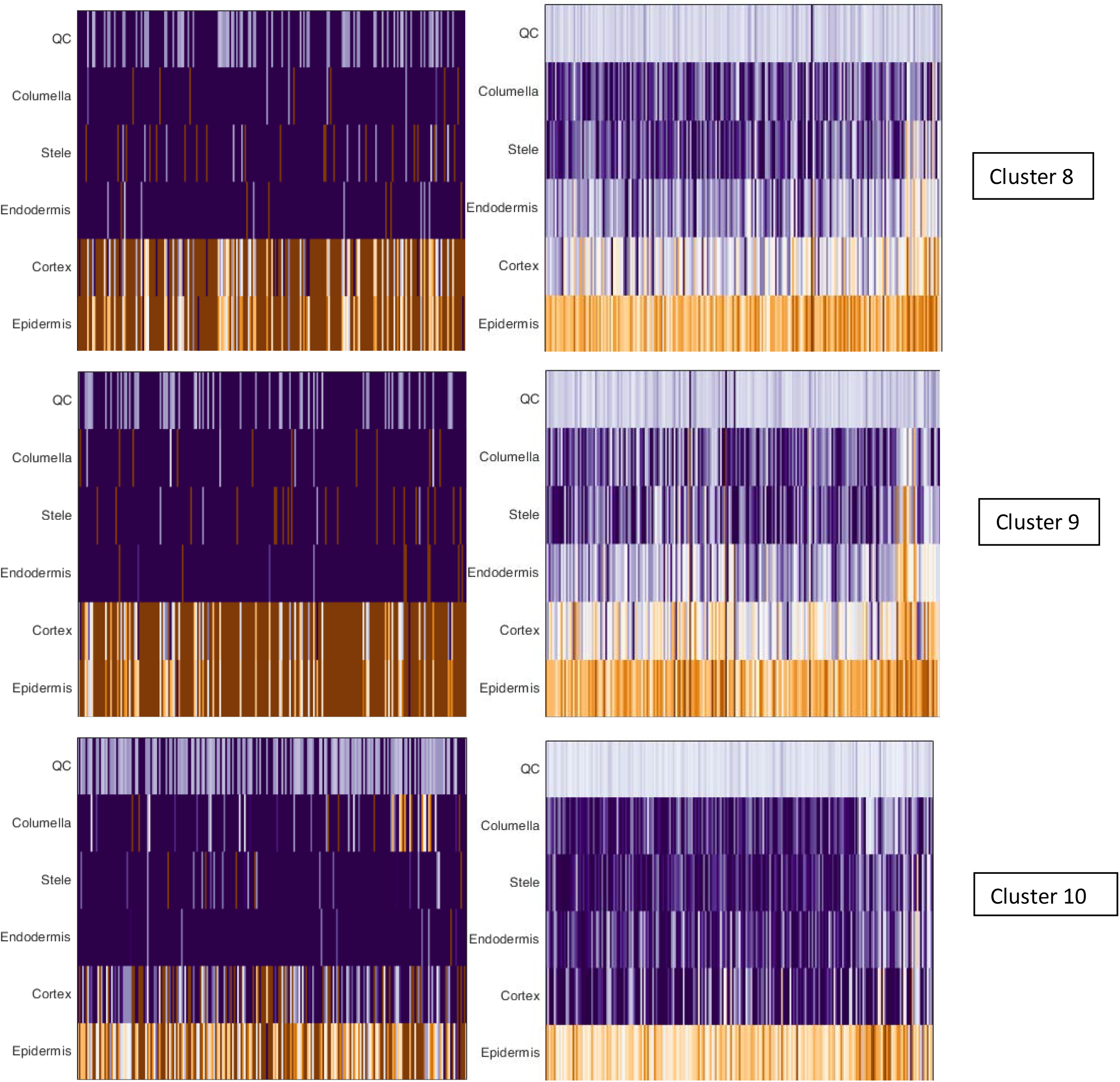
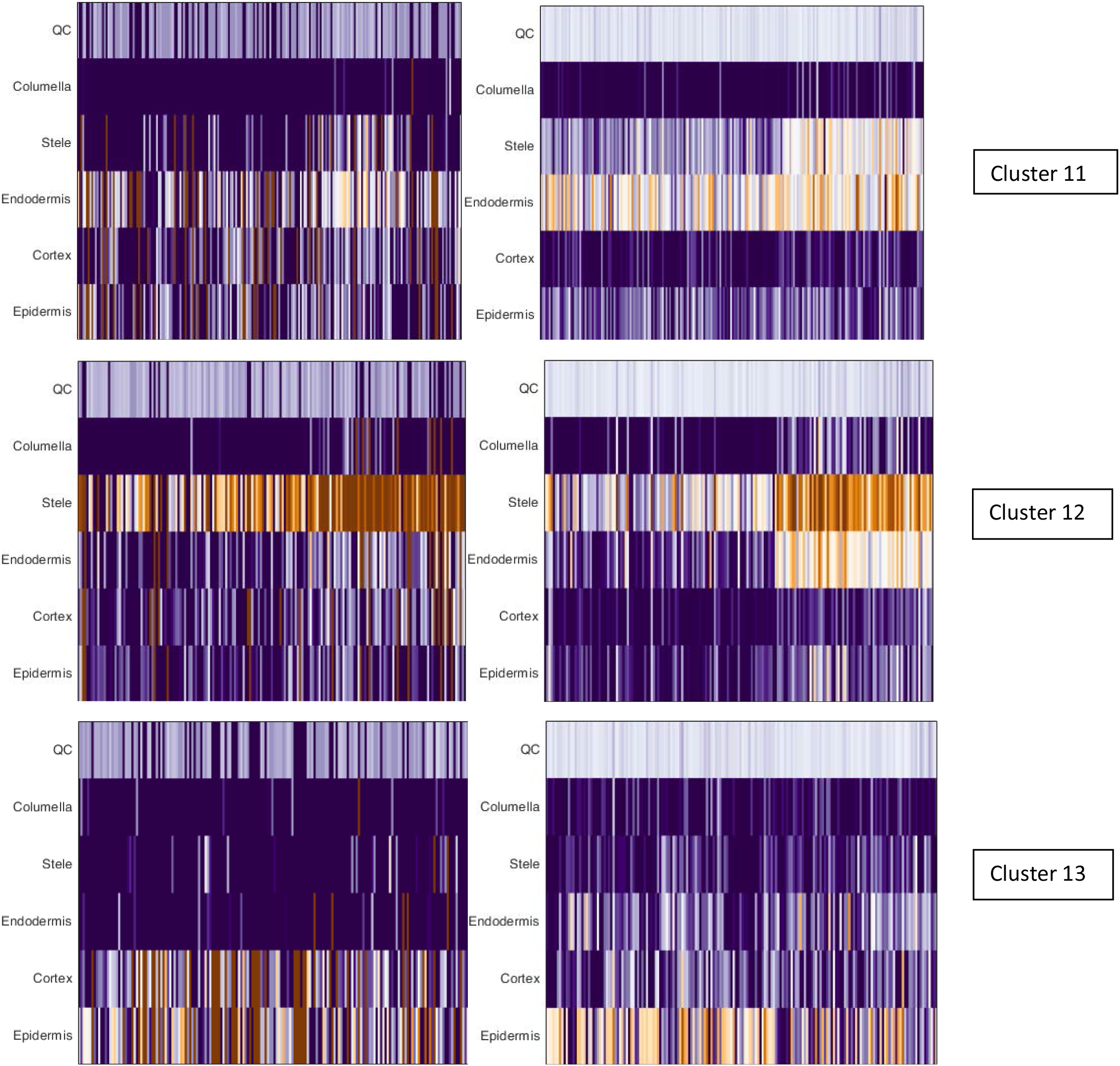
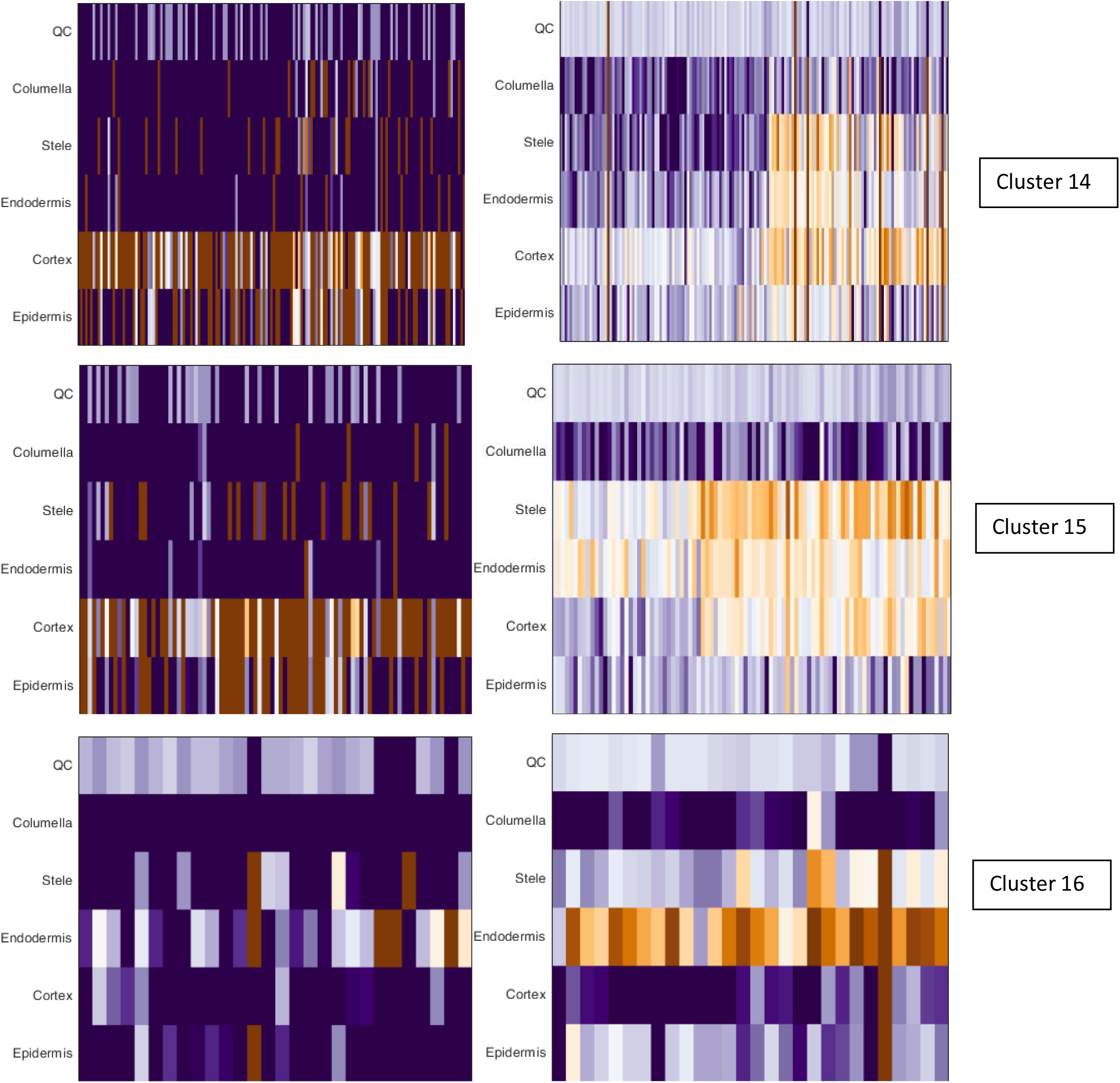
ICI analysis of UMAP defined clusters. Heatmaps show ICI analysis of each cell (columns) for each cluster. The six different cell types represent the six different tissues profiled by either cell sorting or dissection (root cap or columella). The first column of heatmaps represents an ICI analysis with the 25 best markers for each tissue type (stringent set, see Materials and Methods). The second column of heatmaps represent an ICI analysis with the 150 best markers for each tissue type. When the 25-marker analysis gave a weak signal (e.g., cluster 6), the 150-marker analysis was used to narrow down the identification of the cluster in combination with known markers for cell identity (a judgment call given weak quantitative signals). In some rare instances (e.g., cluster 16), the presence of well-documented markers was used to call cell identities over the 150-marker ICI analysis when the stringent 25-marker analysis signal was weak.

**Fig. S3.**
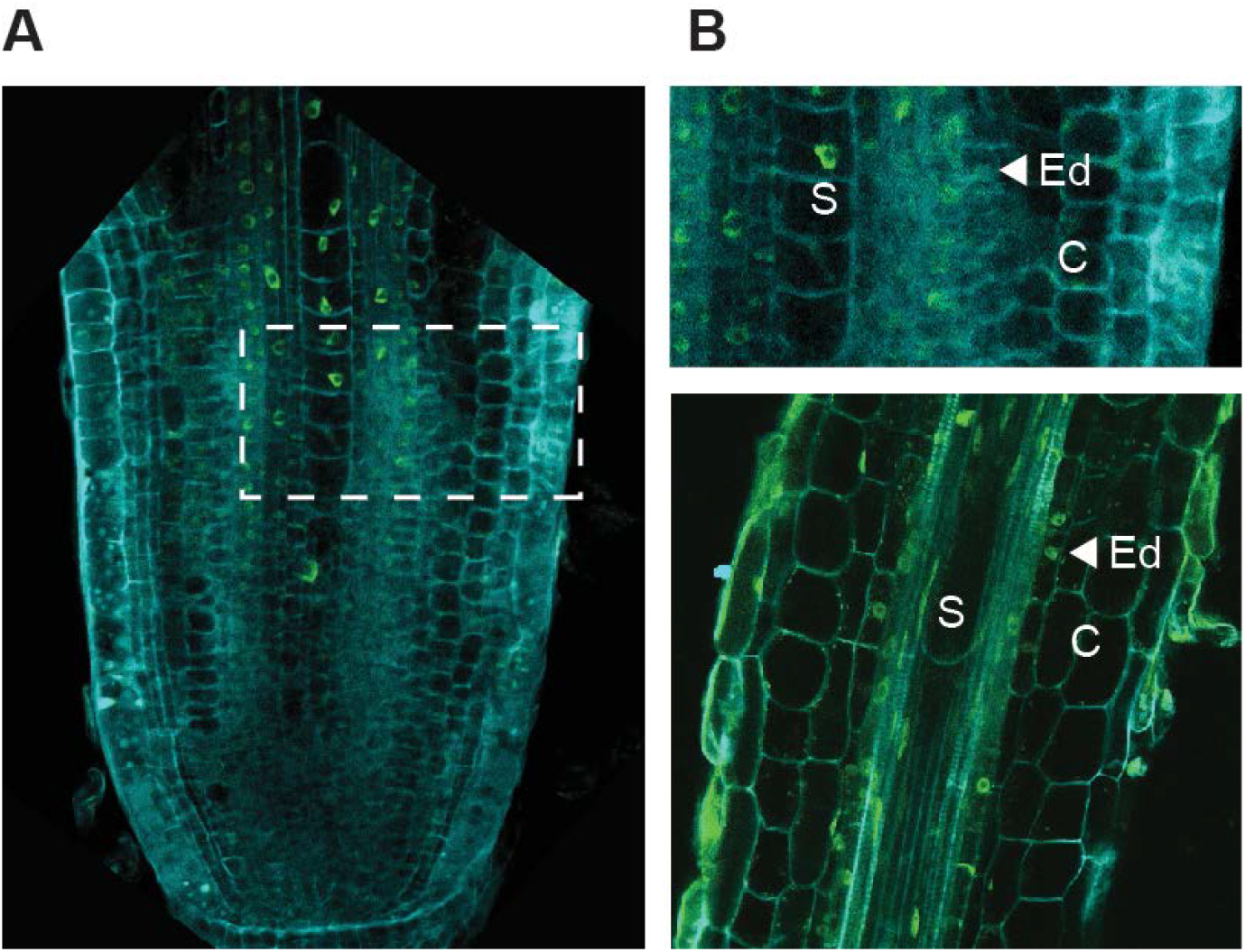
SCR protein is located in the stele in *Setaria viridis* roots. **(A)** *SvSCR* translational reporter showing nuclear localization in stele cells. **(B)** Inset showing enlarged region from dashed box showing endodermis and cortex layers in the root meristem (top), and transition zone (bottom).

**Fig. S4.**
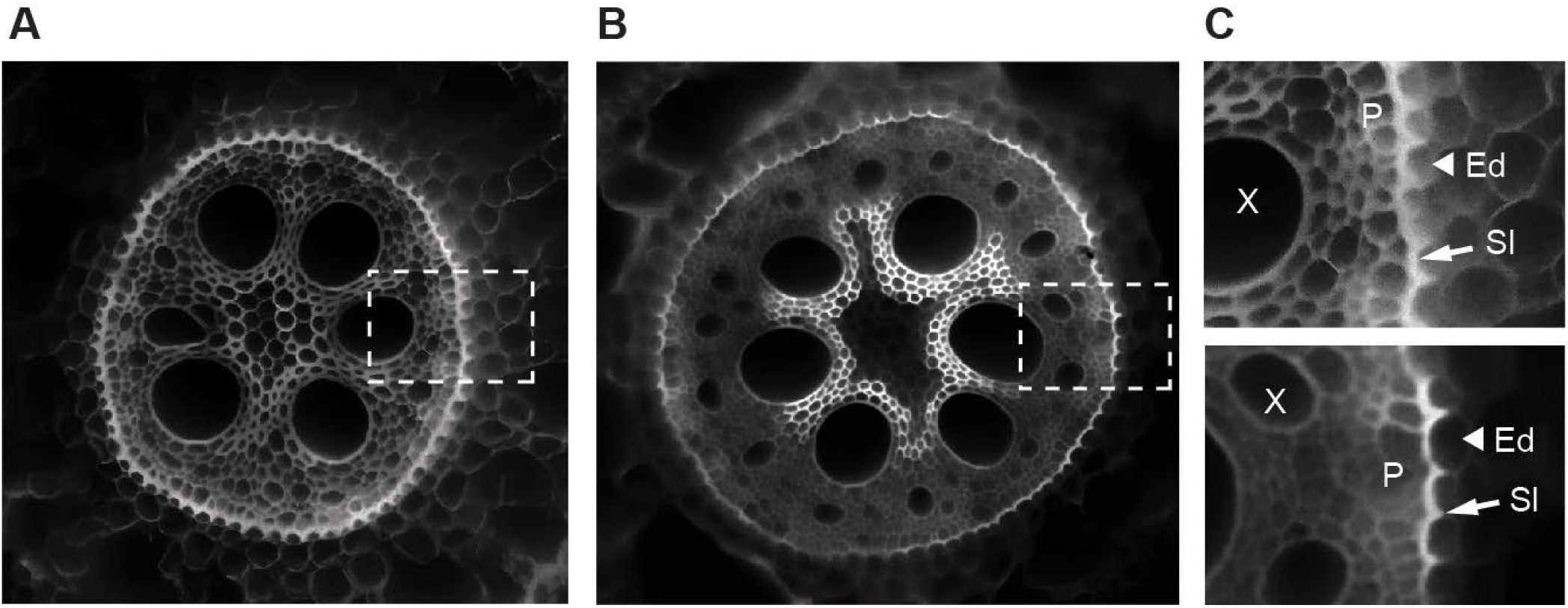
Endodermal identity analysis by suberin staining in wild type and *Zmshr1*/*3* in maize. Wild type and *Zmshr* mutant roots stained with Yellow Fluor 88, which stains the suberin layer (SI) of endodermis (Ed). (A) Cross section of wild type roots showing lens shaped cell with Yellow Fluor88 staining marking an endodermal layer. (B) Cross section of *Zmshr2*/*2-h* mutant also showing the lens-shaped cell layer. (C) Magnification of hatched boxes in C and D, showing wild type (top) and *Zmshr2*/*2-h* (bottom). Pericycle (P) and xylem vessels (X) can also be observed.

**Fig. S5.**
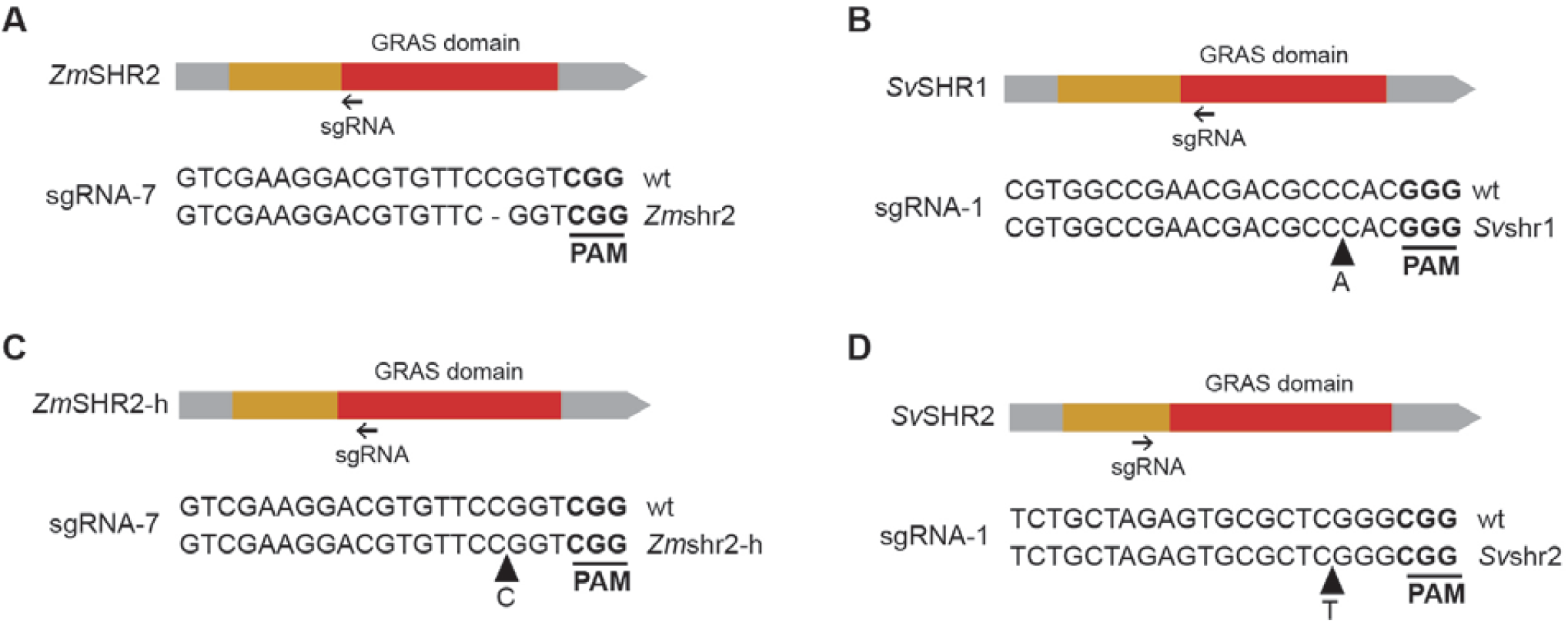
CRSIPR-Cas9 gene editing in ZmSHR and SvSHR genes caused frameshift mutations. (**A & C**). Diagram showing the position of a single-base deletion in *Zm*shr2 and a single-based insertion in *Zm*shr2-h, causing frameshifts. (**B & D**). Diagram showing the position of single-base insertions in *Sv*shr1 and *Sv*shr2. Grey box = UTR region, green = exons, red = SHR GRAS domain, arrows = sgRNAs.

